# Detecting prodromal Alzheimer’s disease with MRI through deep learning

**DOI:** 10.1101/813899

**Authors:** Xinyang Feng, Frank A. Provenzano, Scott A. Small, for the Alzheimer’s Disease Neuroimaging Initiative

**Affiliations:** Department of Biomedical Engineering, Columbia University, New York, NY, 10027, United States; Department of Neurology, Columbia University, New York, NY, 10032, United States; Taub Institute for Research on Alzheimer’s Disease and the Aging Brain, Columbia University, New York, NY, 10032, United States

## Abstract

Deep learning applied to MRI for Alzheimer’s classification is hypothesized to improve if the deep learning model implicates disease’s pathophysiology. The challenge in testing this hypothesis is that large-scale data are required to train this type of model. Here, we overcome this challenge by using a novel data augmentation strategy and show that our MRI-based deep learning model classifies Alzheimer’s dementia with high accuracy. Moreover, a class activation map was found dominated by signal from the hippocampal formation, a site where Alzheimer’s pathophysiology begins. Next, we tested the model’s performance in prodromal Alzheimer’s when patients present with mild cognitive impairment (MCI). We retroactively dichotomized a large cohort of MCI patients who were followed for up to 10 years into those with and without prodromal Alzheimer’s at baseline and used the dementia-derived model to generate individual ‘deep learning MRI’ scores. We compared the two groups on these scores, and on other biomarkers of amyloid pathology, tau pathology, and neurodegeneration. The deep learning MRI scores outperformed nearly all other biomarkers, including—unexpectedly—biomarkers of amyloid or tau pathology, in classifying prodromal disease and in predicting clinical progression. Providing a mechanistic explanation, the deep learning MRI scores were found to be linked to regional tau pathology, through investigations using cross-sectional, longitudinal, premortem and postmortem data. Our findings validate that a disease’s known pathophysiology can improve the design and performance of deep learning models. Moreover, by showing that deep learning can extract useful biomarker information from conventional MRIs, the advantages of this model extend practically, potentially reducing patient burden, risk, and cost.

## INTRODUCTION

Biomarkers can aid in the clinical evaluation of Alzheimer’s disease (AD), and biomarkers currently exist for AD’s three core neuropathologies—amyloid pathology, tau pathology, and neurodegeneration^1,2^. The first two can be estimated from CSF levels of Aβ and tau, or by direct visualization using PET-sensitive radioligands. Neurodegeneration, a term currently used to encompass neuronal or synaptic loss^3^, can be estimated from PET-based measures of parietal cortex metabolism, or MRI-based measurements that reflect the structural integrity of the hippocampal formation.

Deep learning is a subset of machine learning that, in principle, holds promise for MRI-based classification of neurogenerative diseases, including AD^4,5^. Furthermore, while some studies have examined classifying MCI conversion using machine learning frameworks, they have done so using other architectures like SVM^6^, examining only up to 36 months^6-8^, using clinical information in the model^7-9^, and few have examined performance independently against existing biomarkers. We hypothesized that designing a deep learning model that considers AD’s known pathophysiology and anatomy would improve the model’s classification ability. Because ‘cell sickness’ occurs first and foremost in the pathophysiology of AD^3,10,11^, before dramatic neuronal loss, a classifier is predicted to be improved if it is based on alterations in voxel signal intensity rather than on volume shrinkage. Additionally, informed by the brain’s anatomical complexity, particularly the areas whose AD’s pathophysiology targets, a classifier is expected to improve if it is based on 3D than on 2D MRI information.

The challenge with a 3D classifier that depends on voxel signal intensity is that its training is estimated to require an unusually large number of scans from cases and controls, more than is typically available for AD. Having access to large-scale datasets is a common challenge for deep learning in all fields, and strategies have been developed for data augmentation^12^. We deploy a novel data augmentation strategy that is particularly well suited for MRI-only datasets, by including scans acquired from the same patient across multiple visits. By training, validating, and testing the classifier at the level of individual subjects, instead of individual scans, we minimize the potential limitations of this approach, namely data leakage.

We elected not to augment data by traditional methods of image perturbation, like rotating or applying transformations, since structural MRI data have well known preprocessing pipelines to spatially align images. We did not include available clinical information, as studies have done prior^7^, to avoid a model dependent on information that might be sparse or unavailable, as might be the case of clinical evaluation outside of a carefully controlled and harmonized setting, like ADNI.

In the first series of studies, we used this data augmentation strategy to accumulate a large-scale dataset of MRI scans generated from patients with AD dementia and controls, and from which we could test our hypothesis about a deep learning model that uses an intensity-dependent 3D classifier. Confirming our hypothesis, the model, which generates individual ‘deep learning MRI’ scores reflecting AD probability, was found to classify AD dementia with very high accuracy. Moreover, although voxels from the whole brain were included in the model, the most predictive areas turned out to encompass the hippocampal formation. This anatomical profile supports the biological premise of our classification, potentially placing our deep learning MRI scores within the ‘neurodegeneration’ biomarker category.

While these results were encouraging, AD progresses through a prodromal stage before causing dementia, presenting clinically as mild cognitive impairment (MCI)^13^. Only a subset of patients with MCI has prodromal AD, and in contrast to AD dementia, where a clinical evaluation is often sufficient to diagnose the disease, our ability to diagnose prodromal AD when presented with an MCI patient is currently inadequate. With increased awareness and concern over AD, a growing number of MCI patients are presenting to clinicians wanting to know whether they have prodromal AD, and, if so, how quickly they will progress to dementia. Showing that the deep learning algorithm can address the clinical questions that relate to prodromal AD would not only better validate its classification capabilities, but since derived from conventionally-acquired MRI scans, would potentially expand its clinical utility.

Accordingly, in the second series of studies we set out to test how well the deep learning MRI scores, derived from the deep learning model trained on AD dementia, performs in detecting prodromal AD and in predicting time to dementia progression. Additionally, we compared its performance to other biomarkers of amyloid pathology, tau pathology, and neurodegeneration. Based on the premise of deep learning’s classification abilities, we hypothesized that deep learning MRI scores would outperform other MRI-based biomarkers of neurodegeneration. At the same time, given the proposed temporal profile of AD’s neuropathology^14^, we hypothesized that amyloid or tau biomarkers would outperform the deep learning MRI score in classifying prodromal AD. Additionally, we investigated the link of deep learning MRI scores to amyloid and tau pathology, using cross-sectional, longitudinal, premortem and postmortem data, providing mechanistic explanation for the deep learning MRI score.

The diagnostic cutoffs for all AD biomarkers are traditionally derived from patients in the dementia stage, and biomarkers shift over the disease’s progressive course, particularly dynamic during its early stages. Since cutoffs for prodromal AD have not yet been established for any of the biomarkers, the best experimental design with which to test these hypotheses is to clinically follow a large group of MCI patients as they progress to dementia, so that the patients can be retroactively dichotomized into those with and without prodromal AD at baseline. Biomarkers can then be tested to determine which best classifies prodromal AD and which best predicts progression. The challenge with this design is that, based on current estimates, approximately 5 years of clinical follow-up is needed in order to allot sufficient time for the majority of prodromal AD patients to clinically manifest as dementia^15,16^. Here, we were able to implement this experimental design thanks to the Alzheimer’s Disease Neuroimaging Initiative (ADNI), which has been acquiring biomarker data in a large population of MCI patients since 2005 and to test the two hypotheses about which biomarker best classifies prodromal AD, and which predicts progression to dementia.

## RESULTS

### Classifying the dementia stage of Alzheimer’s disease

The deep learning model was trained, validated, and tested on 975 MRI scans repeatedly acquired in patients in the dementia stage of AD, versus 1943 MRI scans repeatedly acquired from healthy controls. In the test set, a ‘deep learning MRI’ score was derived for each scan from the model, with the score reflecting the probability of each scan having AD. A receiver operating characteristic (ROC) analysis revealed that the deep learning MRI scores accurately classified AD dementia vs. healthy controls with an AUROC (area under the receiver operating characteristics curve) of 0.973 (Fig. 1a).

**Figure 1.**
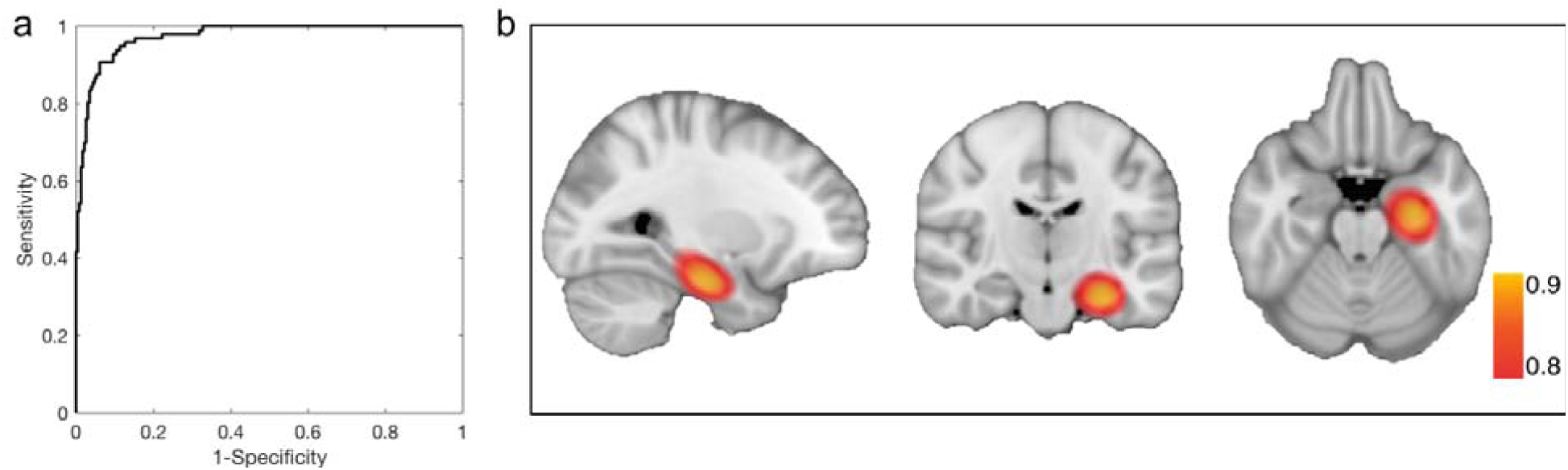
Classifying Alzheimer’s disease in its dementia stage. The ‘receiver operating characteristic’ curve shows that the deep learning MRI score applied to the test set of Alzheimer’s disease (AD) dementia scans vs. healthy controls scans classified AD dementia with high accuracy (panel ‘a’). The class activation map, reflective of the regional contributions to the deep learning MRI scores, localized to the left anterior medial temporal lobe in the vicinity of the entorhinal cortex and hippocampus, where Alzheimer’s pathophysiology begins.

Next, we generated an AD ‘class activation map’ to determine whether the deep learning MRI scores derived from the model were regionally dominated. We find that the deep learning MRI scores are dominated by alterations in voxel signal intensity that localized to anterior medial temporal lobe, in the vicinity of the anterior entorhinal cortex and hippocampus (Fig. 1b). We note that while the class activation map localized to the left more than the right anterior medial temporal lobe, in agreement with previous findings^17-19^, contralateral areas emerged with lowered thresholding.

### Classifying the prodromal stage of Alzheimer’s disease

From ADNI, we identified a cohort of participants who were diagnosed with MCI at baseline and who had a complete set of CSF amyloid and tau biomarkers and structural MRI (N = 582; the inclusionary and exclusionary algorithm is illustrated in Fig. S1). Among these, 205 participants progressed to AD dementia at follow up (‘MCI progression’ group), and thus had prodromal AD at baseline, while 179 participants remained MCI stable for at least 4 years (‘MCI stable’ group) (Fig. 2). The dementia-derived deep learning classifier was used to generate deep learning MRI scores on each individual case.

**Figure 2.**
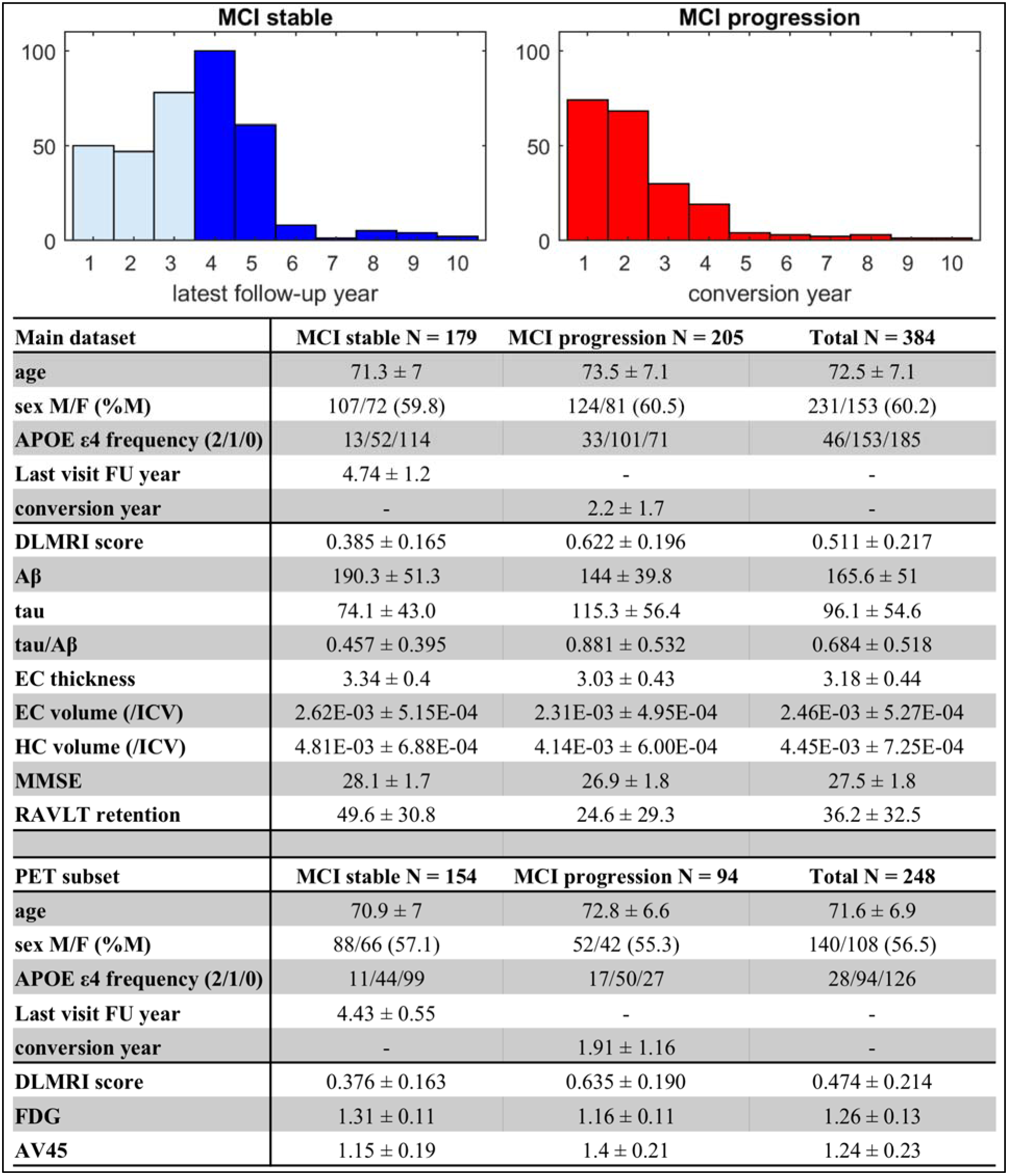
Distribution and demographics of subjects in the ‘mild cognitive impairment’ study. Distribution frequencies of the participants with amnestic mild cognitive impairment (MCI) at baseline, who either remained stable (MCI stable) or progressed to Alzheimer’s dementia (MCI progression), organized by latest follow-up years and conversion years. The dark blue bars indicate participants included in the study. Demographic and baseline biomarker data are listed in the table for the MCI stable and MCI progression groups.

ROC analyses revealed that the deep learning MRI score outperformed all other biomarkers in classifying the MCI-stable from the MCI-progression group (Fig. 3). The AUROC of deep learning MRI score was 0.788 (Accuracy at Youden (ACC)=75%), superior to CSF Aβ (AUROC=0.702 ACC=66.7%, significantly lower than the deep learning MRI score, p=0.0141), CSF tau (AUROC=0.682, ACC=66.4%, p=0.0161), CSF tau/Aβ (AUROC=0.703, ACC=68.5%, p=0.0161); superior to MRI-based measures of hippocampal volume (AUROC=0.733, ACC=67.7%, p=0.0484), entorhinal cortex volume (AUROC=0.64, ACC=62.5%, p=2.01E-6), and entorhinal cortex thickness (AUROC=0.685, ACC=64.1%, p=1.71E-4); and, finally, superior to Mini-Mental State Exam (AUROC=0.648, ACC=63.3%, p=6.70E-5), and to neuropsychological measure most sensitive to the early stages of AD, the RAVLT retention score^20^ (AUROC=0.686, ACC=67.7%, p=2.28E-3).

**Figure 3.**
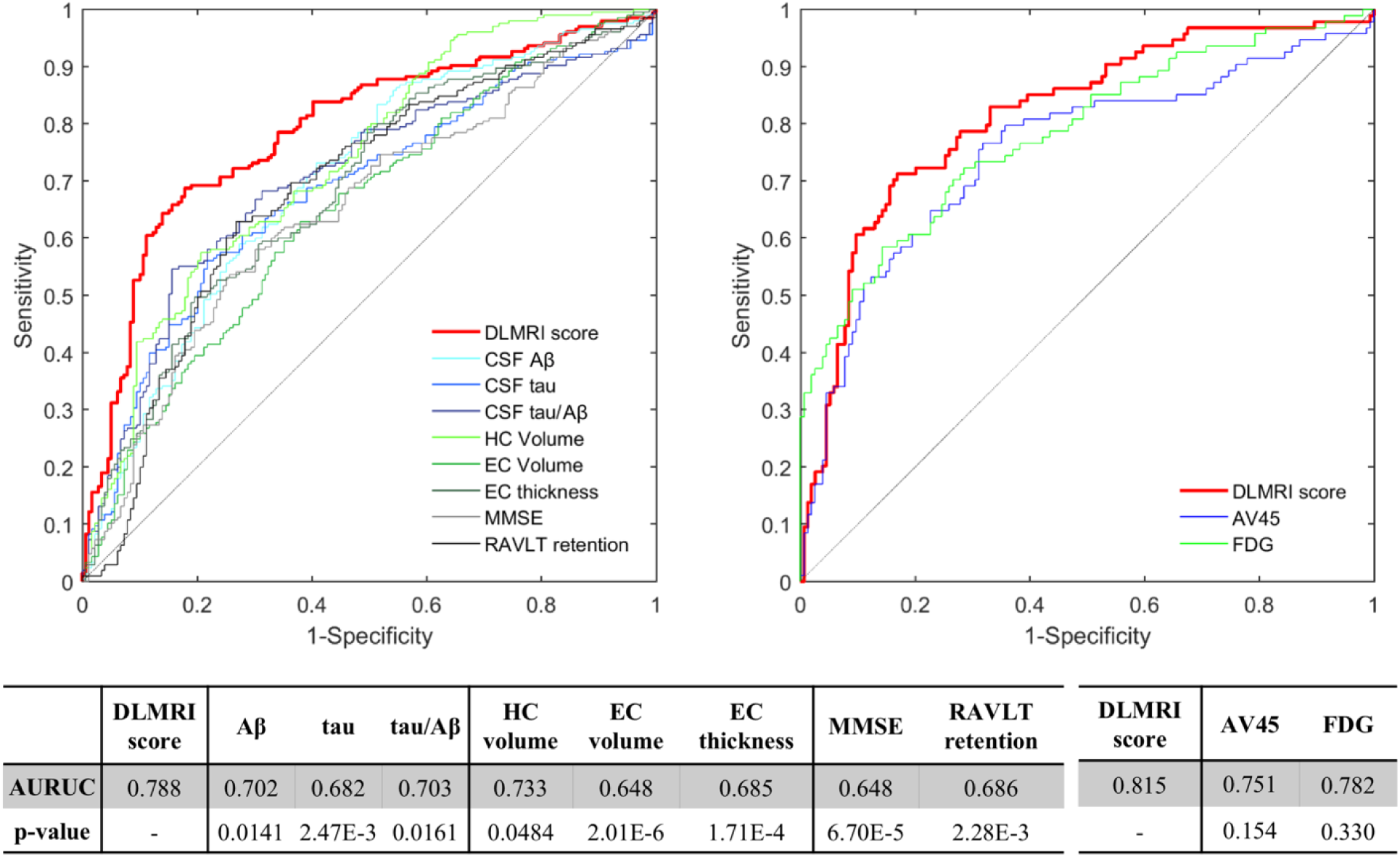
Classifying Alzheimer’s disease in its prodromal stage. By comparing the ‘MCI stable’ to the ‘MCI progression’ groups, ROC curves show that the deep learning MRI (DLMRI) scores were superior in classifying prodromal Alzheimer’s disease (indicated in red). The deep learning MRI scores outperformed (left panel) CSF measures of Aβ, tau, or tau/Aβ; MRI measures of hippocampal (HC) or entorhinal cortex (EC) volume or thickness; clinical measures using the modified mental status exam (MMSE) or the retention of the Rey Auditory Verbal Learning Task (RAVLT) (left panel). In a smaller subset, the deep learning MRI scores (right panel) outperformed PET measures of amyloid using the AV45 radioligand or metabolism using fluorodeoxyglucose (FDG). Specific area under the curve (AUROC) values for each measure, and statistical probability values for each comparison, are shown in the table.

Additionally, the deep learning MRI score was found to outperform or perform as well when tested in a subset of participants in whom additional PET-based biomarkers were available -- FDG-PET that by measuring parietal cortex metabolism is considered a biomarker of neurodegeneration^21^, and AV45-PET, which by using an amyloid radioligand is a biomarker of amyloid pathology^22^. In this subset, the deep learning MRI score classified prodromal AD with an AUROC=0.815 (ACC=78.6%), compared to the AUROC of 0.782 (for PDG-PET (ACC=75.4%) and 0.751 (ACC=71.4%) for amyloid-PET, although the differences were not statistically significant (Fig. 3, bottom panel).

### Predicting progression to Alzheimer’s disease dementia

Survival analyses were performed to determine which biomarker best predicted progression to AD dementia among the MCI groups. Results revealed that compared to other biomarkers, the deep learning MRI score best predicted time to conversion to AD dementia, as illustrated by the survival curves of high and low deep learning MRI scores and tau/Aβ ratios (Fig. 4). The deep learning MRI scores showed better prediction capability (|z|=11.0, p=4.35E-28) than CSF biomarkers of amyloid and tau pathology (Aβ |z|=6.37, p=1.87E-10, tau |z|=5.70, p=1.18E-08, tau/Aβ |z|=5.41, p=6.29E-08); than MRI-based biomarkers of neurodegeneration (hippocampal volume |z|=8.80, p=1.35E-18, entorhinal volume |z|=6.02, p=1.75E-09, entorhinal thickness |z|=7.42, p=1.21E-13); and, than behavioral measures (MMSE |z|=5.72, p=1.07E-08, RAVLT retention |z|=6.88, p=6.12E-12). Similarly, in the subset in whom the additional PET biomarkers were available the deep learning MRI score (|z|=9.04, p=1.40E-19) outperformed or performed as well as FDG-PET (|z|=9.11, p=8.14E-20) and AV45-PET |z|=7.12, p=1.04E-12).

**Figure 4.**
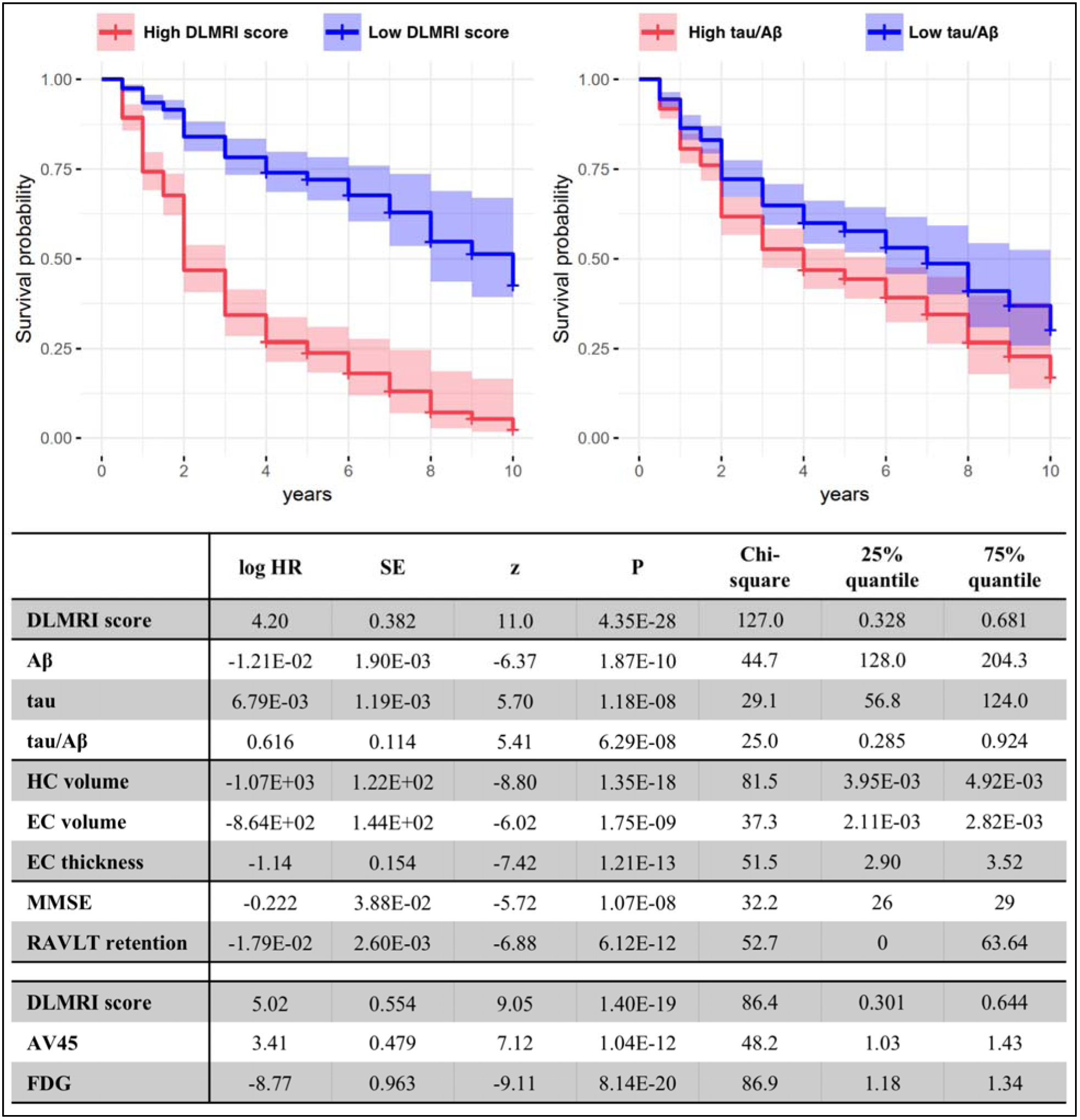
Predicting progression to Alzheimer’s Dementia. Survival analyses were performed comparing the deep learning MRI scores to other measures, and example curves illustrate that the deep learning MRI score (left panel) outperforming the CSF measure of the tau/Aβ ratio (right panel). The high risk (indicated by red) and low risk (indicated by blue) curves were fitted from 75% and 25% percentile of the measures respectively. The shaded area indicates the 95% confidence interval. The deep learning MRI scores outperformed CSF Aβ, tau, or tau/Aβ, MRI-derived measures of hippocampal volume, entorhinal cortex volume, and entorhinal thickness; behavioral measures, Mini-Mental State Exam (MMSE), and RAVLT retention; and, when available, PET measures of amyloid using the AV45 radioligand or metabolism using fluorodeoxyglucose (FDG).

### Correlations with amyloid pathology and tau pathology

Correlational analyses were performed to determine whether the deep learning MRI score was correlated more with amyloid pathology or tau pathology. Cross-sectionally, we found that while the deep learning MRI score showed a stronger correlation with CSF tau (r=0.225, p=9.00E-6), it also correlated with CSF Aβ (r=-0.190, p=1.86E-4). Longitudinally, however, changes in the deep learning MRI scores over time were significantly associated with changes in CSF tau (r=-0.205, p=1.50E-3), but not with changes in CSF Aβ (r=-8.18E-3, p=0.900).

Next, in a subsample with available postmortem data, we correlated the deep learning MRI score with neuropathological evidence of amyloid pathology, as indicated by the Thal staging^23^, or tau pathology indicated by Braak staging^24^. The deep learning MRI scores were found to associate more with tau pathology (with an MRI-autopsy interval below 2 years, Braak staging: r=0.397, p=7.70e-3; Thal staging: r=0.196, p=0.203) (Fig. 5 bottom panel). To further explore the regionality of this relationship, we found that the deep learning MRI score correlated with tau levels mapped by tau-PET, with strong correlations observed with tau pathology in the entorhinal cortex (r=0.449, p=1.66E-15).

**Figure 5.**
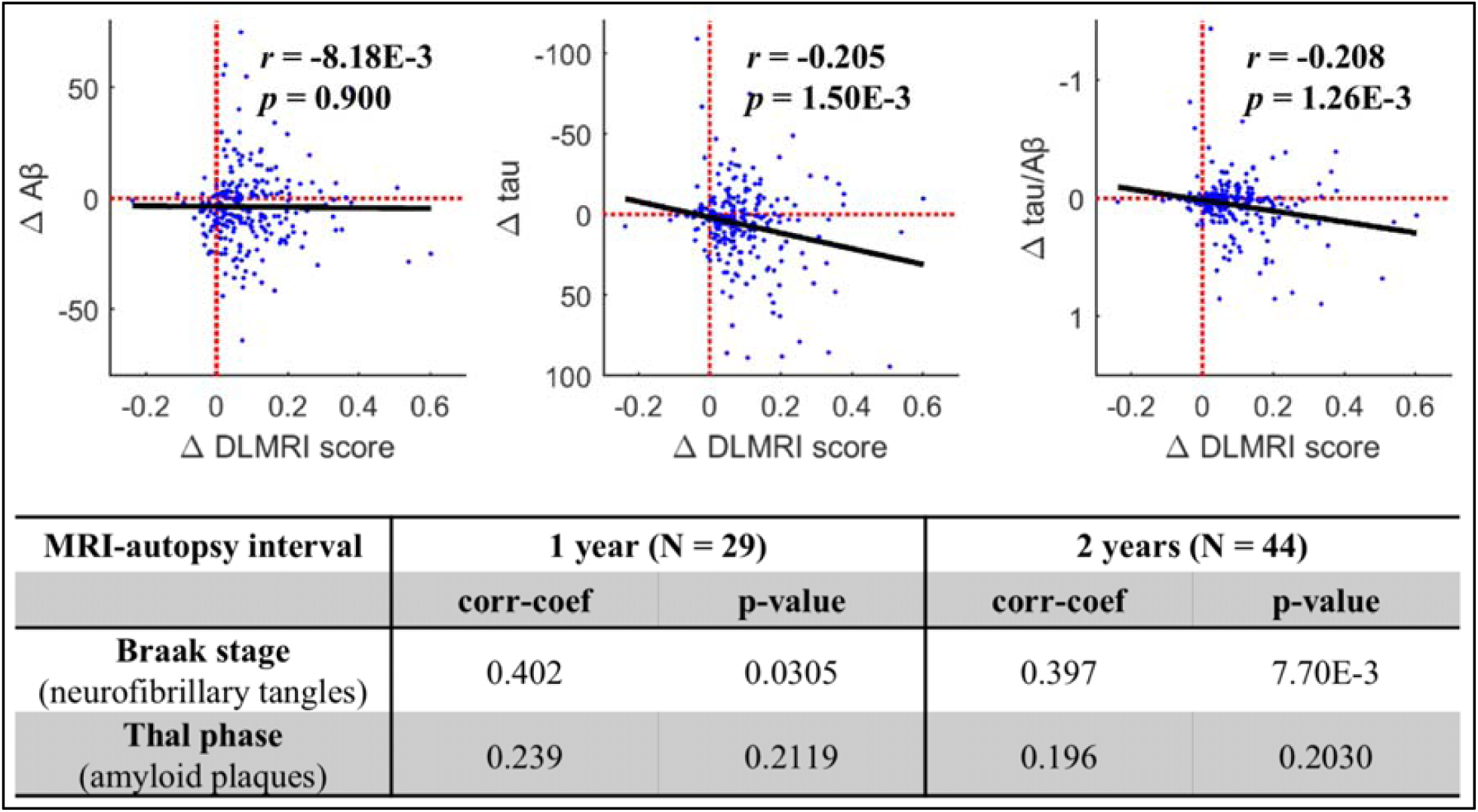
The deep learning MRI score correlates with tau pathology. The scatter plots illustrate the relationship between changes over time in the deep learning MRI scores vs. changes in CSF Aβ (left panel), changes in CSF tau (middle panel) and changes in CSF tau/Aβ (right panel). Each data point indicates one participant’s change of last deep learning MRI score from baseline (ΔDLMRI_last_), plotted against their fitted change in biomarker measures at ΔDLMRI_last_ with the slope estimated from all follow-up visits (see Methods). The black solid lines are the linear fits across participants, showing that changes in the deep learning MRI score is most strongly correlated with changes in tau over time. The table lists the correlations between antemortem deep learning MRI scores to postmortem-derived Braak stage of neurofibrillary tangles and the Thal phase of amyloid plaques, with an MRI-autopsy interval below either 1 year and 2 years, showing that the deep learning MRI scores are most strongly correlated with tau pathology.

## DISCUSSION

The level of performance achieved by our deep learning model in classifying AD dementia supports our hypothesis that a disease’s pathophysiology should be considered when evaluating performance as well as justifying the data augmentation strategy used. Further validating the assumptions, design, and implementation of our model is the fact that, despite incorporating information from the whole brain, the class activation map was dominated by signal in the anterior entorhinal cortex and hippocampus, precisely where AD pathophysiology begins^3,17-19,24^.

Stronger validation of the deep learning model was provided by the second series of studies when the dementia-derived classifier was applied to the prodromal stages of AD. Supporting the first hypothesis of this study, we found that our deep learning MRI scores outperformed other MRI-based measures of neurodegeneration in both classifying prodromal AD and in predicting progression to dementia. Refuting the second hypothesis, we found that the deep learning MRI scores performed at least as well and typically outperformed biomarkers of amyloid and tau pathology.

We do not consider this unexpected finding a challenge to the primacy of amyloid and tau pathology in the neuropathological progression of AD^25^. The deep learning MRI scores were found strongly linked to tau pathology in the entorhinal cortex, a region where AD pathology begins^24^, and its superior performance likely reflects this sensitivity. It is possible, therefore, that tau-PET would outperform the deep learning MRI score and other biomarkers. ADNI, however, has only begun acquiring tau-PET in 2015, and there is currently insufficient data to test this prediction in our experimental design. Future analyses from ADNI and other long-term PET studies will be able to test this prediction.

The observation that the deep learning MRI scores outperformed biomarkers of amyloid and tau pathology in predicting time to dementia is less surprising. As a biomarker of neurodegeneration, this finding agrees with prior studies^26^ and with the current model for the temporal sequence of AD’s neuropathology^25^. Since in this scheme neurodegeneration occurs last, accurate biomarkers of it are more proximal to the development of dementia.

The strength of our prodromal AD study is that by relying on progression to AD dementia as a way to retroactively identify patients with prodromal AD, we overcame the limitation that precise biomarker cutoffs for prodromal AD have not yet been established. We designed the analysis based on prior studies that suggest that the majority of MCI patients with prodromal AD will progress to dementia within 4-5 years^15^, an assumption confirmed in our study. Furthermore, approximately half of the MCI cohort ended up having prodromal AD, which agrees with previous approximations^27^. Still, a potential weakness of our study is the possibility that a minority of patients in the stable MCI category are harboring prodromal AD at baseline. The number of misclassified patients is likely to be low^27^, and so this potential imprecision would not be expected to significantly alter our results. Tracking stable MCI patients for longer periods might address this concern, but would in fact raise a new one: when tracking patients for a decade or more, particularly given the high incidence of AD in older populations, some are expected to develop AD *de novo* after the baseline evaluation. We can conclude that our findings and their conclusions are beyond reproach for a 5-year time window after initial evaluation, a clinically meaningful epoch for both patients and clinicians.

Our study provides the proof-of-principle that imaging-based deep learning models that are examined in concert with a disease’s pathophysiology will yield a highly accurate model and improve performance in prognosticating disease. Showing that deep learning can enhance the utility of MRI in prodromal AD is the more important clinical implication of this study. Ordering “neuroimaging studies”^28^ is the current standard of care when evaluating a patient with MCI suspected of having AD, most typically the conventional MRIs from which the deep learning MRI scores were derived. The rationale for this recommendation and its routine clinical implementation is not to ‘rule in’ AD, but rather to exclude other non-neurodegenerative causes of dementia, such as strokes, bleeds, and tumors. Deep learning algorithms that can extract useful information for the purposes of prodromal AD detection, from conventional MRIs that have in any case been acquired, has the additional advantages of reducing patient burden and cost incurred by lumbar punctures, injection of radioactive ligands, or other additional testing.

## METHODS

### Participants in the Alzheimer’s disease dementia study

All data were obtained from ADNI, a multi-site observational study, which were acquired in accordance with each site’s respective Institutional Review Board, including obtaining written consent acquired from each participant. We included 2918 scans (N_healthy control_ = 1943, N_AD_ = 975) from 626 subjects as training set, 382 scans (N_healthy control_ = 251, N_AD_ = 131) from 80 subjects as validation set, and 325 scans (N_healthy control_ = 229, N_AD_ = 96) from 80 subjects as test set.

Our data augmentation method of using scans from multiple visits of the same participant requires dealing with two problems: data leakage and disease progression. Data leakage is the problem of including different scans from the same participant in the training and test set, the trained model might make the prediction by matching the subject instead of extracting disease relevant patterns. In this study, the training, validation and test sets were partitioned at subject level to ensure non-overlapping subjects. Disease progression is the problem that the diagnosis status of subjects might change during follow-up visits, and the diagnosis at scan time might be different from the baseline label. In this study, we labeled all the scans with their cross-sectional diagnosis at scan time.

### Participants in the ‘Mild Cognitive Impairment’ study

From ADNI we identified a cohort of participants who were diagnosed with MCI at baseline and who had a complete set of CSF amyloid and tau biomarkers and structural MRI (N = 582; the inclusionary and exclusionary algorithm is illustrated in Fig. S1). Among these, 205 participants progressed to AD dementia at follow up (‘MCI progression’ group), and 179 participants remained MCI stable for at least 4 years (‘MCI stable’ group). The time distribution and demographics of these two groups are shown in Fig. 2.

### The deep learning MRI score

The deep learning model used in this study is a three-dimensional convolutional neural network (3D CNN) model with five convolutional stages and one fully connected layer with sigmoid output^5^. Each convolutional stage consists of two convolutional layers with rectified linear unit (ReLU) activation function, a batch normalization operation and a max pooling layer. The model was optimized using ADAM method with cross-entropy loss, using a learning rate of 2e-5 determined through grid search. The model was trained on the brain-extracted T1-weighted structural MRI scans from the ADNI cohort to classify patients in the dementia stage of AD versus healthy control subjects. To evaluate the regional contribution to AD classification, we generated a 3D class activation map, which visualizes the predictive regions in deep learning classification models^29,30^.

We applied the model trained to classify AD dementia versus healthy controls to the baseline scans of patients diagnosed with MCI. The continuous output from the model is reflective of the progressive structural patterns of AD pathology. We refer to it as a ‘deep learning MRI’ score. All analyses were performed using this score.

### Amyloid and Tau Biomarkers

#### CSF biomarkers

CSF tau levels, reflective of neurofibrillary tangle, and CSF Aβ levels, reflective of amyloid pathology, were included in the analysis^31^. Additionally, the tau/Aβ ratio, which has been shown to best capture AD^32^, was also included^33^. CSF was acquired at individual ADNI sites in accordance to the ADNI acquisition protocols and analyzed as previously described^33^. The median values provided by ADNI were used.

#### PET measures

In a subset of participants (N_MCI-progression_ = 94, N_MCI-stable_ = 154), amyloid pathology was also estimated with PET, mapping amyloid burden with the amyloid-binding radioligand AV45. The composite AV45-PET score provided by ADNI^34^ was used in the analyses, which is based on the average AV45 SUVR (standard uptake value ratio) of the frontal, anterior cingulate, precuneus, and parietal cortex relative to the cerebellum^35^.

### Neurodegeneration Biomarkers

#### MRI morphometry

FreeSurfer 6.0^36,37^ was used to segment the structural MRI scans and derive regional morphometric measures. Hippocampal (HC) volume, entorhinal cortex (EC) volume, and entorhinal cortex thickness were used as structural integrity measures of the hippocampal formation. Hippocampal and entorhinal cortex volume were normalized by the intra-cranial volume (ICV).

#### PET measures

In a subset of participants (N_MCI-progression_ = 94, N_MCI-stable_ = 154), neurodegeneration was also estimated with PET using fluorodeoxyglucose (FDG). The composite FDG score provided by ADNI^34^ was used in the analyses, which is based on the average FDG uptake of angular, temporal, and posterior cingulate^21^.

### Additional Measures

#### Behavioral and neuropsychological measures

The MMSE (Mini-Mental State Examination) score and RAVLT (Rey Auditory Verbal Learning Test) retention scores were used in the analysis. The RAVLT retention score measures the number of delayed recalled words divided by the number of words learned in the last learning trial (trial 5) and has been found to be one of the most sensitive to AD^20^.

#### Neuropathology

Among subjects with postmortem neuropathology data, 44 cases were identified who had an MRI within two years prior to death, and 29 cases were identified who had MRI within one year prior to death. DLMRI scores were derived from the last antemortem MRI scans in this cohort. An association was investigated between DLMRI score and the neuropathologically-derived Braak stage, which reflects neurofibrillary tangles^24^, and the Thal phase, which reflects amyloid plaques^23^.

#### Tau-PET

ADNI began acquiring PET scan using the AV1451 radioligand, which binds neurofibrillary tangles^38^, in the late phase of ADNI2 and resumed in ADNI3. Due to the smaller number of subjects with available longitudinal tau-PET data or follow-up visits, cross-sectional analyses on these subjects (N = 296) using the regional AV1451 retention levels provided by ADNI^34^ were performed.

### Statistical analysis

#### ROC analysis

A receiver operating characteristic (ROC) analysis was used to determine the accuracy of the deep learning MRI score in prodromal AD classification, i.e. MCI-stable and MCI-progression classification, using standardized residuals controlling for age, sex, and APOE ε4 frequency with linear regression. The DeLong test^39^ was used to test for the significance of the differences in the AUROCs (area under the ROC curve) between DLMRI score and other measures using the pROC R package^40^.

#### Survival analysis

Cox proportional hazards regression models were fit to examine the association between each baseline measure and time to conversion to AD dementia from MCI, controlling for age, sex, and APOE ε4 frequency, using the survival R package^41^. MCI-stable participants are included in the models as censored data with the last visit as the censored point. The high-risk and low-risk survival curves were generated with the 75% percentile and 25% percentile of the observed measures, respectively.

#### Longitudinal analysis

The longitudinal association between DLMRI score and CSF biomarkers was studied by examining the deviation from baseline measurements for each participant over time. From the ‘MCI-progression’ and ‘MCI-stable’ group, we further identified participants that had at least one follow-up of both MRI and CSF, and collapsed them into a group for longitudinal analysis (n=238). The changes in either CSF biomarker or DLMRI score of all follow-up visits from baseline were used to estimate the slope β of the change in tau (Δtau), Aβ (ΔAβ), and tau/Aβ ratio (Δtau/Aβ) versus the change in DLMRI score (ΔDLMRI) for each participant using linear regression through the origin. Each participant was represented by the point based on the last follow-up visit’s ΔDLMRI_last_ (x-coordinate) and the fitted change βΔDLMRI_last_ (y-coordinate) of the respective measure. The last follow-up visit was used to anchor the representation of the participant in order to reflect the full follow-up. A correlation analysis was performed across participants. A linear regression model was fit across participants and illustrated.

#### Correlational analysis

A partial correlation was performed between baseline DLMRI score and CSF biomarkers, regional tau-PET measures, controlling for age, sex, and APOE ε4 frequency. As the Braak stage of neurofibrillary tangles and the Thal phase of amyloid plagues are both rank ordinal measures, we correlated the DLMRI score with the neuropathological measures using Spearman correlation.

## ACKNOWLEDGEMENTS

Data collection and sharing for this project was funded by the Alzheimer’s Disease Neuroimaging Initiative (ADNI) (National Institutes of Health Grant U01 AG024904) and DOD ADNI (Department of Defense award number W81XWH-12-2-0012). ADNI is funded by the National Institute on Aging, the National Institute of Biomedical Imaging and Bioengineering, and through generous contributions from the following: AbbVie, Alzheimer’s Association; Alzheimer’s Drug Discovery Foundation; Araclon Biotech; BioClinica, Inc.; Biogen; Bristol-Myers Squibb Company; CereSpir, Inc.; Cogstate; Eisai Inc.; Elan Pharmaceuticals, Inc.; Eli Lilly and Company; EuroImmun; F. Hoffmann-La Roche Ltd and its affiliated company Genentech, Inc.; Fujirebio; GE Healthcare; IXICO Ltd.; Janssen Alzheimer Immunotherapy Research & Development, LLC.; Johnson & Johnson Pharmaceutical Research & Development LLC.; Lumosity; Lundbeck; Merck & Co., Inc.; Meso Scale Diagnostics, LLC.; NeuroRx Research; Neurotrack Technologies; Novartis Pharmaceuticals Corporation; Pfizer Inc.; Piramal Imaging; Servier; Takeda Pharmaceutical Company; and Transition Therapeutics. The Canadian Institutes of Health Research is providing funds to support ADNI clinical sites in Canada. Private sector contributions are facilitated by the Foundation for the National Institutes of Health (www.fnih.org). The grantee organization is the Northern California Institute for Research and Education, and the study is coordinated by the Alzheimer’s Therapeutic Research Institute at the University of Southern California. ADNI data are disseminated by the Laboratory for Neuro Imaging at the University of Southern California.

## AUTHORS’ CONTRIBUTIONS

XF, FAP, SAS contributed to literature search, figures, study design, data interpretation, and writing. XF contributed to data collection and data analysis.

## FIGURE LEGENDS

**Figure S1.**
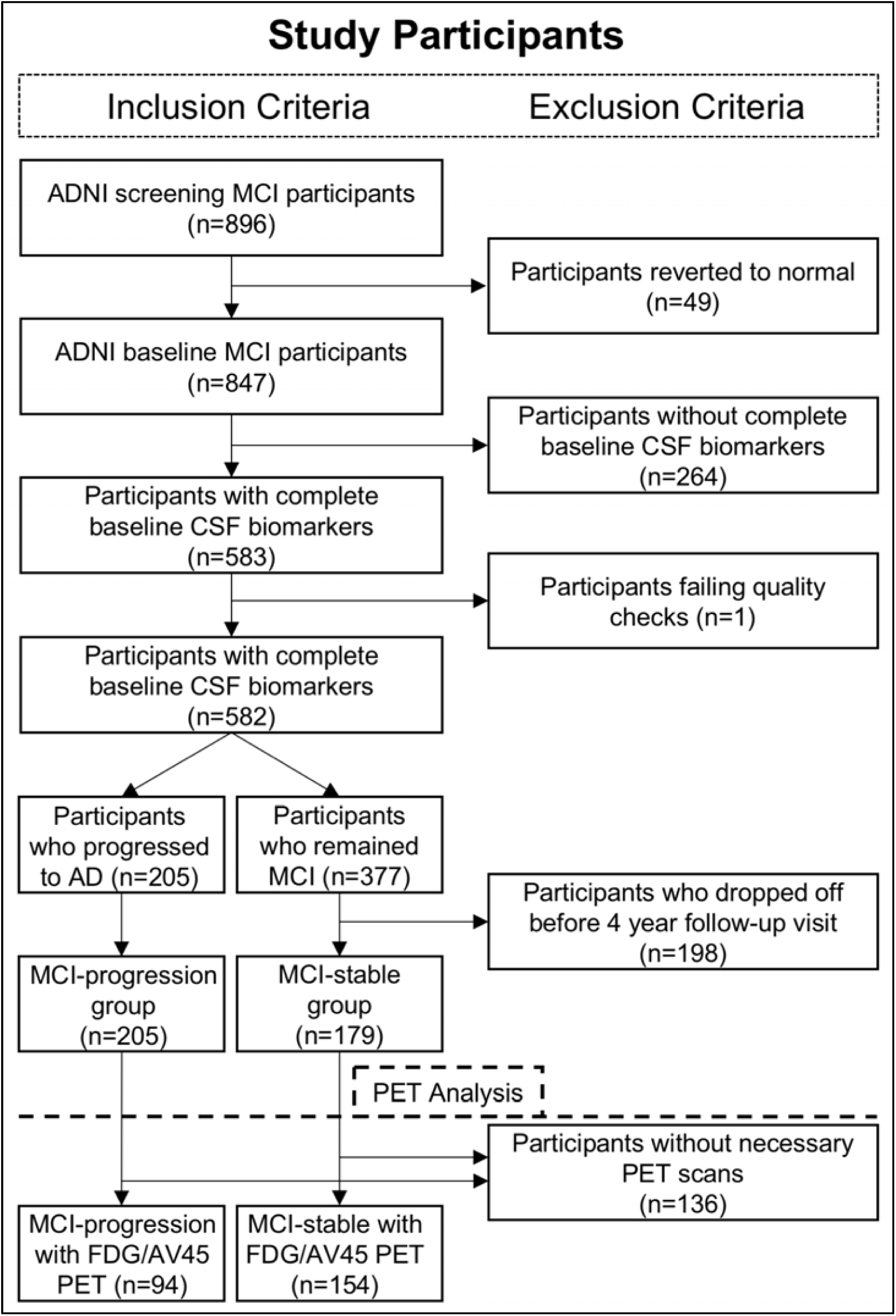
Participant selection flow-chart.

